# The protein domains of vertebrate species in which selection is more effective have greater intrinsic structural disorder

**DOI:** 10.1101/2020.10.15.341313

**Authors:** Catherine Weibel, Jennifer E James, Sara M Willis, Paul G Nelson, Joanna Masel

## Abstract

The effectiveness of selection varies among species. It is often estimated by means of an “effective population size” based on neutral polymorphism, but this is confounded in complex ways with demography. The strength of codon bias more directly pertains to how well adaptation at many sites can be maintained in the face of deleterious mutations, but past metrics that compare codon bias across species are confounded by among-species variation in %GC content and/or amino acid composition. Here we propose a new Codon Adaptation Index of Species (CAIS) that corrects for both confounders. Unlike previous metrics, CAIS yields the expected relationship with adult vertebrate body mass. As an example of the use of CAIS, we ask whether protein domains evolve lower intrinsic structural disorder (ISD) when present in more exquisitely adapted species, as expected given that ISD is higher in eukaryotic proteomes than prokaryotic proteomes. Using phylogenetically corrected linear models, we find, contrary to expectations, that the ISD of a given protein domain evolves to be higher when in well-adapted species. This effect is stronger in young protein domains but is also present in ancient domains.

## Introduction

Species differ from each other in many ways, including mating system, ploidy, spatial distribution, life history, size, lifespan, and population size. These differences have population genetic implications, such that the process of adaptation is more efficient in some species than others. Difficulties in measuring differences in the effectiveness of selection among species currently impede our ability to discover the systematic influence of selection effectiveness on phenotypes.

The mutation-selection-drift model describes how the effectiveness of selection depends on a species’“effective” population size, *N_e_* (Ohta 1973). A population with a smaller *N_e_* has a harder time purging deleterious mutations (Kimura, 1962; Ohta, 1972, 1992). A useful way to operationalize the effectiveness of selection is to consider a one locus, two allele model. Over a long period of time, and in the absence of mutation bias, the effectiveness of selection can then be captured as the ratio of the frequencies of fixed deleterious allele: fixed beneficial allele states, ranging from 1:1 (ineffective selection such that mutation bias sets the ratio) to just over 0:1 (highly effective selection). This ratio can be calculated as the ratio of the probability of fixation of deleterious mutations to the counterfixation of their beneficial alternatives (King and Masel 2007). In a Wright-Fisher model, this ratio depends on the product *sN* (Figure 1).

**Figure 1:**
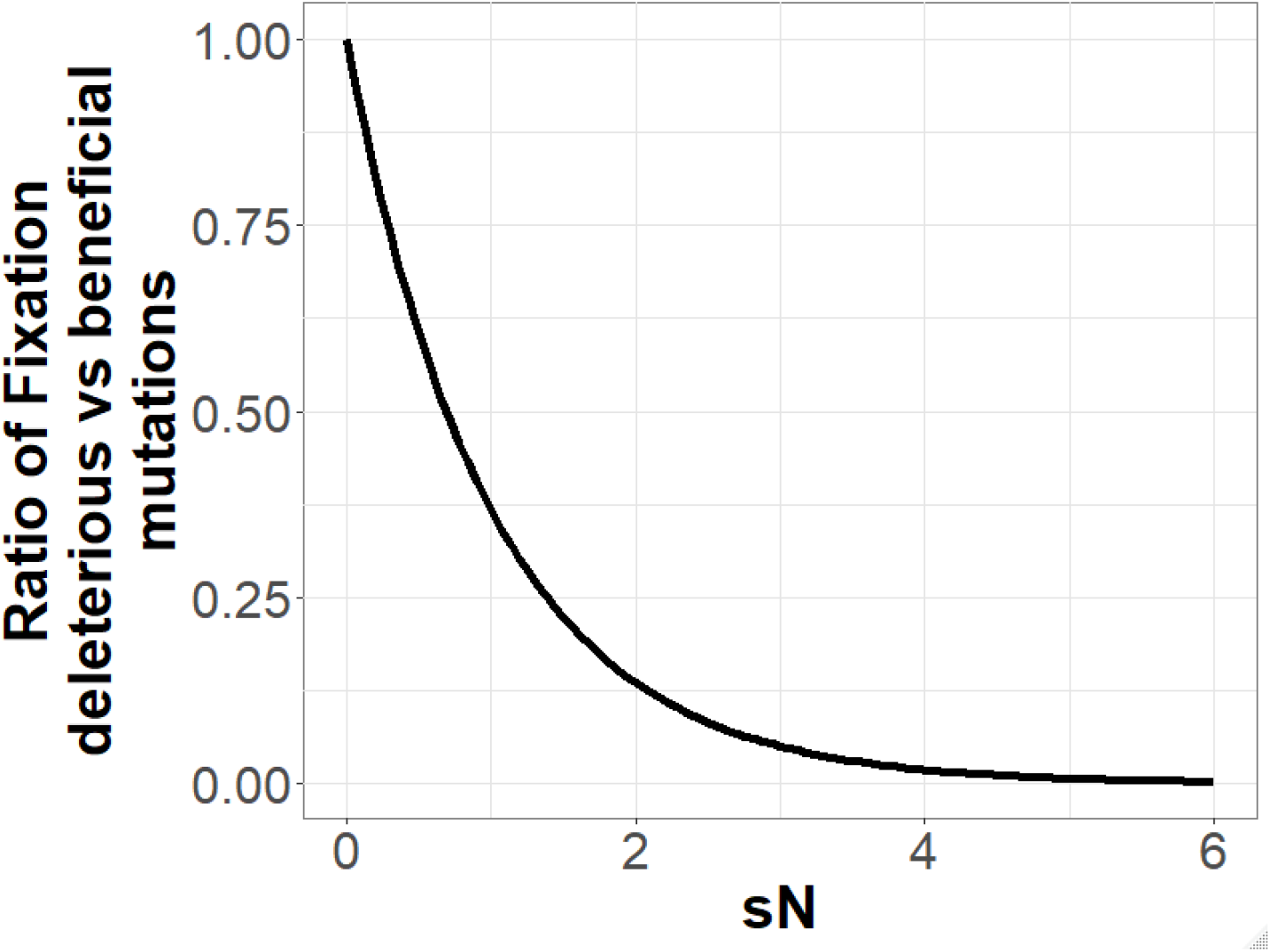
The effectiveness of selection, calculated as the long-term ratio of time spent in fixed deleterious: fixed beneficial allele states given symmetric mutation rates, is a function of the product *sN*. Assuming a diploid Wright-Fisher population with 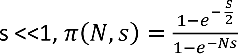, and the y-axis is calculated as *π(N, −s)/π(N,s)*. In the x-axis, *s* is held constant at a value of 0.001 and *N* is varied. Results for other plausible values of *s* are superimposable.

In general, *N_e_* is defined as the value of *N* of an idealized Wright-Fisher population in which a property of interest matches that of a given, real population. The same population can therefore have different values of *N_e_* for different properties of interest. The most commonly used properties concern levels of neutral polymorphism, and the most common method for estimating *N_e_* (Lynch and Conery 2003) is thus to divide a measure of putatively neutral (often synonymous) polymorphism by that species’ mutation rate (Charlesworth 2009). This measure of *N_e_* is only available for species that have both polymorphism data and accurate mutation rate estimates, restricting its use. Worse, it is not a robust statistic (Galtier and Rousselle 2020). In the absence of a clear species definition, polymorphism is sometimes calculated across too broad a range of genomes, substantially inflating *N_e_* (Daubin and Moran 2004).

In any case, the value of *N_e_* important for our purposes is not with respect to neutral polymorphism, but rather with respect to the probability of fixation of slightly deleterious mutations, and hence the degree to which highly exquisite adaptation can be maintained in the face of deleterious mutations with low s (Ohta 1973). To compare species, we therefore focus on the degree of selective preference among synonymous codons as a more direct alternative to assess how effective selection is at the molecular level in a given species (Akashi 1996; Subramanian 2008; Galtier et al. 2018). Synonymous mutations are often subject to weak selection for factors such as translational speed and accuracy (Venetianer 2012), to a degree that varies among genes within a species (Sharp and Li 1986; Sharp et al. 2010). The Effective Number of Codons (ENC) (Wright 1990; Novembre 2002; Fuglsang 2004; Fuglsang 2008; Hershberg and Petrov 2008) and the Codon Adaptation Index (CAI) (Sharp and Li 1986) are common metrics to quantify codon bias.

More highly adapted species will have effective selection on codon bias for a higher proportion of their genes. However, exploiting metrics of codon bias to compare codon bias among species, rather than among genes of the same species, raises new issues. In particular, GC content becomes a confounding factor. Genomic GC content is a major driver of codon usage difference among species, through the degree of GC-biased gene conversion and mutational bias with respect to GC (Eyre-Walker et al. 2002; Hershberg and Petrov 2009; Forcelloni and Giansanti 2020).

The original formulation of ENC quantifies how far the codon usage of a sequence departs from equal usage of synonymous codons (Wright 1990). This creates a complex relationship with GC content, which is not easily disentangled (Fuglsang 2008). Fortunately, ENC been modified to correct for differences in nucleotide composition (Novembre, 2002), allowing us to correct for differences in %GC content among species (Supplementary Figures 1A,1B).

The CAI takes the average of Relative Synonymous Codon Usage (RSCU) scores, which quantify how often a codon is used relative to the codon that is most frequently used to encode that amino acid in that species. The CAI is then normalized by maximum synonymous codon usage (RSCU) values, to attempt to control for differences in amino acid composition across the reference set. To compare species, it has been suggested that only highly expressed reference genes be used (Sharp and Li 1986). Unlike ENC, CAI has not previously been modified to control for GC content (Sharp and Li 1986; Labella et al. 2019; Novoa et al. 2019), making it a measure of codon bias rather than codon adaptation. I.e., the more that GC content departs from 50%, the greater the codon bias will be, even in the absence of selection. This issue is exacerbated if CAI scores are calculated not just for highly expressed reference genes, but across entire proteomes, yielding a substantial dependence on GC content (Supplementary Figures 1C,1D).

Quantifying codon adaptation among species, rather than among genes of the same species, might be confounded not just with the biases that shape GC content but also with factors affecting amino acid composition. Species that make more use of an amino acid for which there is stronger selection among codons would have higher codon bias, even if each amino acid, considered on its own, had identical codon bias irrespective of which species it is in. Neither ENC (Fuglsang 2004; Fuglsang 2008) nor the CAI (Sharp and Li 1986) adequately control for differences in amino acid composition when applied across species. Despite claims to the contrary (Wright 1990), this problem is not easy to fix for ENC (Fuglsang 2004; Fuglsang 2008).

A still more serious problem is that CAI’s normalization term, if applied proteome-wide, “drives the bus” in the wrong direction. The exquisiteness of selection appears on the denominator of RSCU scores (see equation 4, Supplementary Figure 2A). Paradoxically, this can make more exquisitely adapted species have lower rather than higher species-level CAI scores. CAI and RSCUs have been inappropriately used as metrics of codon adaptation across species in a handful of publications, adding to the confusion (Jansen et al. 2003; Labella et al. 2019).

Here we develop a new codon adaptation metric that quantifies the effect of selection across the annotated proteome of a species, corrected for both genomic GC and amino acid composition. Proteome-wide metrics have become more broadly accessible given the proliferation of complete genomes, and allow the effectiveness of selection to be estimated without needing to consider demographic history, mutation rate, gene expression level or reference genes. Controlling ENC for amino acid content in a mathematically sound way is difficult (Fuglsang 2008), so we instead build on the CAI framework for our new Codon Adaptation Index of Species (CAIS).

As an example of the use of CAIS, we go on to investigate protein intrinsic structural disorder (ISD). ISD is more abundant in eukaryotic than prokaryotic proteins (Ahrens et al. 2017; Basile et al. 2019), suggesting that low ISD might be favored by more effective selection (Liberles et al. 2012; Ahrens et al. 2017). This difference in structural disorder between eukaryotes and prokaryotes is strongest in the regions between annotated domains, both in abundance and degree of disorder, but the same difference is also visible in protein domains (Basile et al. 2019).

However, it is possible that the difference between prokaryotes and eukaryotes reflects which protein sequences are present, rather than how a single protein sequence evolves differently in different species as a function of the effectiveness of selection in that species. We focus on the latter as a cleaner indication of how descent with modification varies among species as a function of the effectiveness of selection in that species. To do so, we focus on protein domains, whose homology is well-annotated, and which are a more fundamental evolutionary unit than genes (Bornberg-Bauer et al. 2005; P. Bagowski et al. 2010).

Here we develop a new metric of the degree of codon adaptation in a species, one that controls for variation among species in both GC content and amino acid content. We use it to investigate whether the same Pfam protein domain will evolve higher or lower ISD when present in a species in which selection is more effective.

## New Approaches: Codon Adaptation Index of Species (CAIS)

We developed a new metric, the Codon Adaptation Index of Species (CAIS), which calculates the degree of departure from the synonymous codon usage that is predicted by genomic GC content. As a result, CAIS is uncorrelated with total genomic GC Content, even with phylogenetic correction (p value >0.1, Supplementary Figures 1E,1F).

Like the Codon Adaptation Index (CAI) (Sharp and Li 1986), the CAIS is a geometric mean of codon scores, with higher values indicating preferred codons. Because different species have different amino acid compositions, and because some amino acids might have stronger codon preferences, we calculate the CAIS with respect to a standardized amino acid composition, rather than with respect to the actual amino acid composition of the species. Unlike the CAI and ENC, we include stop codons, whose usage can also be biased (Brown et al. 1990; Drabkin and RajBhandary 1998; Southworth et al. 2018). We describe the construction of the CAIS metric in the Methods.

## Results

### Species with larger body mass have smaller CAIS

We consider how three different metrics of codon usage predict body mass in vertebrates. We expect species with larger body size to have smaller effective population size, and thus less codon adaptation (Doyle et al. 2015). Observed correlations between species properties can be due to phylogenetic confounding, a form of pseudoreplication. In all the following analyses, we therefore control for phylogenetic non-independence using Phylogenetic Independent Contrasts (PIC) (Felsenstein,1985).

The CAI, if taken at face value, would paradoxically suggest that larger species have more effective selection (Figure 2A). This is because the behavior of the CAI is driven by the normalization term on its denominator, rather than by its numerator (See Supplementary Figure 2). This problem is removed in the CAIS, which yields the expected result that species with more codon adaptation according to the CAIS tend to be smaller (Figure 2B).

**Figure 2:**
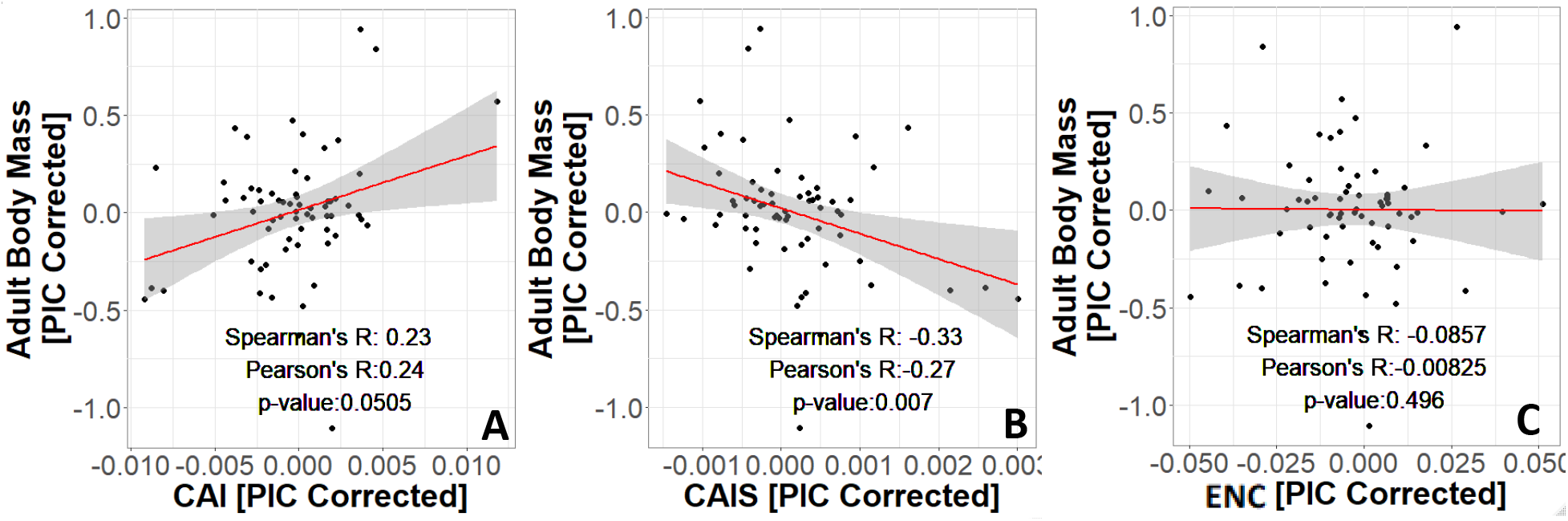
CAIS reflects the expected relationship between effectiveness of selection and body size, while CAI and ENC do not. Body size data are from PanTHERIA database, originally in log_10_(mass) in grams prior to PIC correction. Data are shown for 62 species in common between PANTHERIA and our own dataset of 118 vertebrate species that have both “Complete” genome sequence available for calculating %GC and TimeTree divergence dates. P-values shown are for Spearman’s correlation. Red line shows unweighted lm(y~x) with grey region as 95% confidence interval.

In contrast, the ENC is independent of adult vertebrate body mass, consistent with past reports that effective population size does not predict codon usage in mammals (Figure 2C) (Kessler and Dean 2014). Note that a high ENC value means more codons are being used in the genome of the given species, so that the given species is less codon adapted, while a high CAIS value means that a species is more codon adapted.

This difference in results is surprising because ENC and CAIS are conceptually similar (see Methods). One difference is that CAIS corrects for amino acid composition differences across the dataset while ENC does not (see Methods). However, when we remove the amino acid composition correction from CAIS, we retain the relationship that smaller species tend to have more codon adaptation (Supplementary Figure 3), ruling this out as the cause of their different behaviors. The other key difference between ENC and CAIS is that CAIS is linear in observed codon frequencies (equation 7), while ENC has a quadratic term (equation 12). The quadratic term in ENC may magnify the differences in more extreme deviations of observed frequencies from the GC-informed expected frequencies than CAIS, potentially resulting in the loss of some information in the process.

Given the body size results, we advocate for the use of CAIS as a preferred metric of species’ effectiveness of selection. However, we obtain similar results for ISD when using ENC instead of CAIS; these are shown in the supplement.

### Better adapted species have higher protein disorder

We next consider whether the same homologous domains have lower ISD when found in a more exquisitely adapted species (as assessed by CAIS). ISD, representing a protein’s conformational entropy, can be predicted from amino acid sequence alone using the IUPred program (Dosztányi et al. 2005; Mészáros et al. 2018). While high structural disorder modulates aggregation, it also impedes the efficacy of protein folding which has the potential to impact protein function (Liberles et al. 2012; Macossay-Castillo et al. 2019). Therefore, we might expect proteins to have lower ISD when found in more exquisitely adapted species, in line with the proteome-wide differences between eukaryotes and prokaryotes (Ahrens et al. 2017; Basile et al. 2019).

However, results might be different when tracking the same protein domains than when making proteome-wide comparisons. Species vary both in terms of which protein domains they contain and how many copies of each are present. Higher ISD in eukaryotes might therefore be due to their genomes containing a larger number of high-ISD proteins, rather than because the same domains have higher ISD when in a eukaryote.

To control for the different pfam contents of different species’ genomes, we use a linear mixed model, with a fixed effect on ISD for each species, while controlling for Pfam domain identity as a random effect. We then ask whether those fixed species effects on ISD are correlated with CAIS.

**Figure 3:**
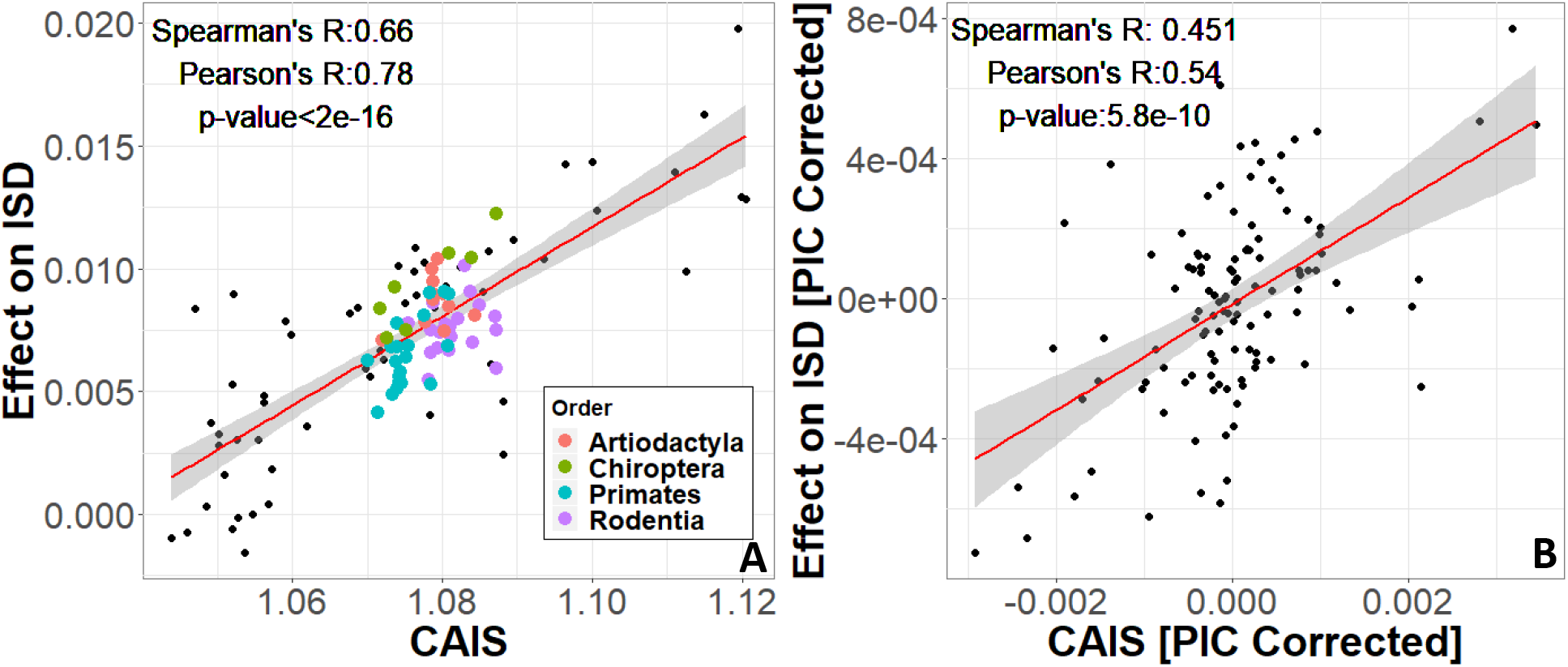
Protein domains have higher ISD when found in more exquisitely adapted species. A) The most common Orders are shown in color; the correlation within each is in the same positive direction as the overall correlation. Each datapoint is one of 118 vertebrate species with “Complete” intergenic genomic sequence available (allowing for %GC correction) and TimeTree divergence dates (allowing for PIC correction). P-values shown are for Spearman’s correlation. Red line shows unweighted lm(y~x) with grey region as 95% confidence interval. Surprisingly, more exquisitely adapted species have more disordered protein domains (Figure 3A). With phylogenetic correction, the strong positive correlation between CAIS and the Species Effect on ISD weakens slightly but is still highly significant (p = 5.8e-10). Results are similar using ENC instead of CAIS (Supplementary Figure 4).

**Figure 4:**
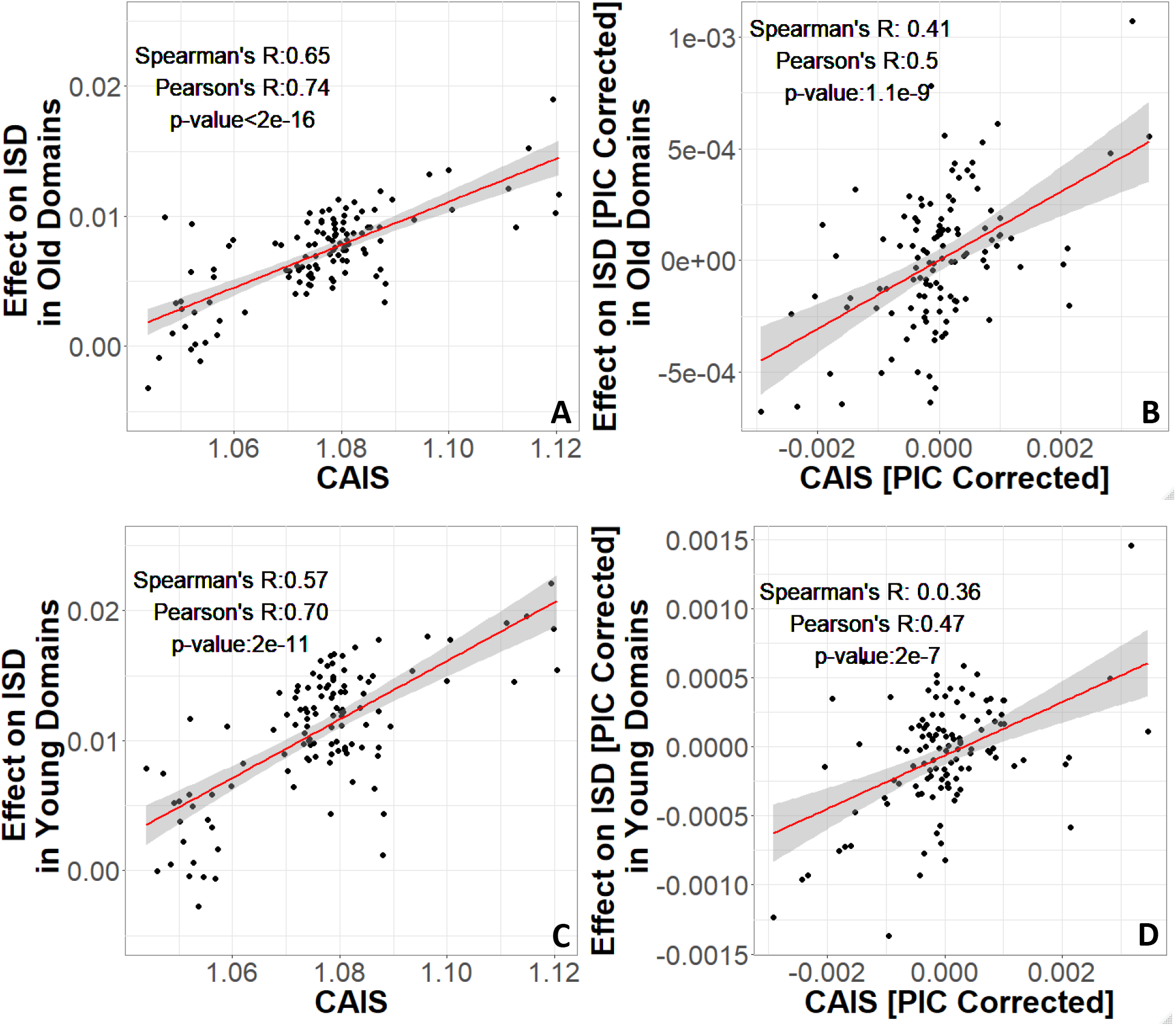
More exquisitely adapted species have higher ISD in both ancient and recent protein domains. In analysis, protein domains that emerged prior to LECA are identified as “old”, and protein domains that emerged after the divergence of animals and fungi from plants and found in vertebrates are identified as “young”. Age assignments are taken from (James et al. 2020). “Effects” on ISD shown on the y-axis are fixed effects of species identity in our linear mixed model. The same n=118 datapoints are shown as in Figure 3. P-values shown are for Spearman’s correlation. Red line shows lm(y~x),with grey region as 95% confidence interval, using a weighted model for non-PIC-corrected figures and unweighted for PIC-corrected figures. Weighted models make more accurate the comparison between young/old domains.

James et al. (2020), when looking just at animal-specific domains, saw higher ISD in young domains. However, there was no such trend among the different ages of old domains (all predating animals). We therefore hypothesize that selection in favor of high ISD might be strongest in young domains, which use more primitive methods to avoid aggregation (Foy et al. 2019; Bertram and Masel 2020). To test this, we analyze two subsets of our data: those that emerged prior to the last eukaryotic common ancestor (LECA), here referred to as “old” protein domains, and “young” protein domains that emerged after the divergence of animals and fungi from plants. Young and old domains both show a trend of increasing disorder with species’ adaptedness (Figure 4). Because PIC analysis makes the units incommensurable, we quantitatively compare the slopes of non-PIC-corrected weighted regressions. As expected, the slope is stronger among young protein domains than among old domains (0.223 +/− 0.021 versus 0.161 +/− 0.014, respectively; units of proportion ISD/CAIS), although it is striking how much even ancient domains prefer higher ISD. See Supplementary Figure 5 for confirmation of these results using ENC.

## Discussion

Here we propose CAIS as a new metric to quantify how species differ in the effectiveness of selection. CAIS corrects codon bias both for total genomic GC content and for amino acid composition, to extract a measure of codon adaptation. Unlike the ENC, CAIS is sensitive enough to show the expected relationship with adult vertebrate body mass. As an illustration of how the CAIS can be used, we estimated the effect of vertebrate species on ISD, while controlling for Pfam identity as a random effect in a linear model. Using phylogenetically controlled linear models, we find that the same Pfam domain tends to be more disordered when found in a well-adapted species (i.e. a species with a higher CAIS). This is true for both ancient and recent protein domains.

The CAIS controls for GC content, which is the product of many different processes. There might be selection on individual nucleotide substitutions, hypothesized to favor higher %GC (Long et al. 2018). Likely more potent are genome-wide forces of mutation bias and gene conversion. Gene conversion is intrinsically biased toward GC and can resemble the effects of selection (Romiguier and Roux 2017). We note however that the magnitude of gene conversion can itself be the target of selection (Gossmann et al. 2012). Mutational biases can either increase or decrease %GC content, and can also be the target of indirect selection on %GC (Smith and Eyre-Walker 2001; Hershberg and Petrov 2009; Hildebrand et al. 2010; Novoa et al. 2019; Forcelloni and Giansanti 2020).

We control CAIS for genomic %GC, which in the vertebrates we study is dominated by intergenic %GC (Galtier et al. 2018). This captures the effects of both mutational biases and GC-biased gene conversion, and so excludes these forces from influencing CAIS. CAIS thus captures the extent of adaptation in codon bias, including translational speed, accuracy, and any intrinsic preference for GC over AT that is specific to coding regions. These remaining codon-adaptive factors do not create a statistically significant correlation between CAIS and GC (Supplementary Figures 1E,1F). This agrees with studies of random ORFs in *E. coli,* where fitness was driven more by amino acid composition than %GC content, after controlling for the intrinsic correlation between the two (Kosinski et al. 2020). Note that we have not ruled out selection for higher %GC in ways that are general rather than restricted to coding regions, whether in shaping mutational biases and the extent of gene conversion, or even at the single nucleotide level in a manner shared between coding regions and intergenic regions. Because the amino acids that promote high ISD are intrinsically GC-rich (Ángyán et al. 2012), it is only appropriate to ask about the evolution of ISD among species after we control for %GC, as Supplementary Figures 1E and 1F show we have done successfully.

Note that if a species were to experience a sudden reduction in population size, e.g. due to habitat loss, leading to less effective selection, it would take some time for CAIS to adjust. CAIS represents a relatively long-term historical pattern of adaptation. The timescales setting neutral polymorphism based *N_e_* are likely shorter (Gossmann et al. 2012).

ISD might be subject to different evolutionary processes at short vs long timescales. Here we found that at relatively short timescales, evolution via selective descent with modification favors high ISD in vertebrates. High ISD is also favored during the gene birth process (McLysaght and Hurst 2016; Wilson et al. 2017; Foy et al. 2019; James et al. 2020; Kosinski et al. 2020). Our results showing a preference for high ISD are surprising given that prokaryotes are more exquisitely adapted than eukaryotes at the molecular level (Liberles et al. 2012; Ahrens et al. 2017), yet have lower ISD (Ahrens et al. 2017; Basile et al. 2019). The obvious reason for the apparent discrepancy is that prokaryotes and eukaryotes contain different protein-coding sequences. In agreement with this view, animal domains that were born longer ago, as evidenced by being found in species across the tree of life today, have lower ISD (James et al. 2020).

Given the influence of ISD on gene birth, all apparent discrepancies would be resolved if higher ISD sequences were differentially lost altogether, while retained sequences are under shortterm selection for higher ISD. Selection might thus work differently on two different timescales, via differential retention on longer timescales vs selective descent with modification on shorter timescales. This hypothesis of different processes on different timescales is consistent with toy models of protein evolution, in which there is a short-term gain to hydrophilicity, but one which impedes the likelihood of eventually finding a stable fold that balances protein stability versus aggregation propensity (Bertram and Masel 2020). This hypothesis is also consistent with our finding that the youngest sequences, which have done the least to obtain a stable fold, are under the strongest short-term selection for higher ISD.

Here we developed a new metric of species adaptedness at the codon level, one capable of quantifying degrees of codon adaptation even among vertebrates. We chose vertebrates partly due to the abundance of suitable data, and partly as a stringent test group, given past studies suggesting limited evidence for codon adaptation. We restricted our analysis to only the best annotated genomes, in part to ensure the quality of intergenic %GC estimates, and in part limited by the feasibility of running massive mixed linear models with six million data points. The phylogenetic tree is well resolved for vertebrate species, with an overrepresentation of mammalian species. Despite the focus on vertebrates and resultant quantitatively tiny differences among species, we see a remarkably strong signal for a subtle codon adaptation effect across closely related species, to the point where we can comfortably detect the ISD signal across subsets of domains.

Our new CAIS metric can be estimated for far more species than an effective population size based on neutral polymorphism, and more directly quantifies how species vary in their exquisiteness of adaptation. We expect CAIS to have many uses, as a new tool for exploring nearly neutral theory.

## Methods

### Species and domains

Pfam sequences and IUPRED2 estimates of Intrinsic Structural Disorder (ISD) predictions were taken from (James et al. 2020), who studied species marked as “Complete” in the GOLD database, with divergence dates available in TimeTree (Kumar et al. 2017). (James et al. 2020) applied a variety of quality controls to exclude contaminants from the set of Pfams and assign accurate dates of Pfam emergence. Pfams that emerged prior to LECA are identified as “old”, and pfams that emerged after the divergence of animals and fungi from plants are identified as “young”, as annotated by (James et al. 2020).

We restricted our analysis to vertebrates, the most species-rich phylogenetic group in James et al. (2020), yielding 170 species. Of the 118 vertebrates in our dataset, 62 species had body size data available through the PanTHERIA database (Jones et al. 2009). We transformed the data by taking log10(body size (g)).

### GC Content

We calculated total GC content (intergenic and genic) during a scan of all six reading frames across genic and intergenic sequences available from NCBI with access dates between May and July 2019 (code available at https://github.com/cweibel2018/More_adapted_species_have_higher_SD.git). Of the 170 vertebrates, 118 had annotated intergenic sequences within NCBI, so we restricted the dataset further to keep only the 118 species for which total GC content was available.

### Codon Adaptation Index

(Sharp and Li 1986) quantified codon bias through the Codon Adaptation Index (CAI), a normalized geometric mean of synonymous codon usage bias across sites, excluding stop and start codons. We modify this to calculate CAI including stop and start codons. While usually used to compare genes within a species, among-species comparisons can be made using a reference set of genes that are highly expressed in yeast genome (Sharp and Li 1986). Each codon *i* is assigned a Relative Synonymous Codon Usage value:

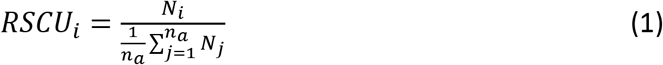

where *N_i_* denotes the number of times that codon *i* is used, and the denominator sums over all *n_a_* codons that code for that specific amino acid. RSCU values are normalized to produce a relative adaptiveness values *w_i_* for each codon, relative to the best adapted codon for that amino acid:

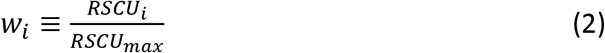

(Sharp and Li 1986) describe the relative adaptiveness values as a control for amino acid composition.

Let *L* be the number of codons across all protein-coding sequences considered. Then

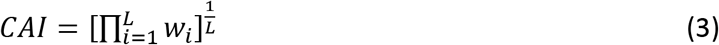

To understand the effects of normalization, it is useful to rewrite this as:

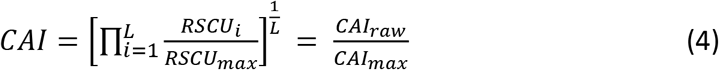

where *CAI_raw_* is the geometric sum of the “unnormalized” or observed synonymous codon usages, and *CAI_max_* is the maximum possible observed CAI given the observed codon frequencies.

### Codon Adaptation Index of Species (CAIS)

#### Controlling for GC bias in Synonymous Codon Usage

Consider a species in which the proportion of the genome that is G or C =*g.* With no bias between C vs. G, nor between A vs. T, nor patterns beyond the overall composition taken one nucleotide at a time, the expected probability of seeing codon *i* in a random sequence is

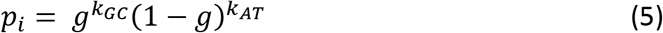

where *k_GC_ + k_AT_* = 3 total positions in codon *i*. The expected probability that amino acid *a* is encoded by codon *i* is

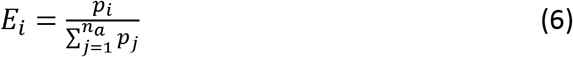

The Relative Synonymous Codon Usage (RSCU) value used by the CAI measures the degree to which a codon’s relative frequency differs from the null expectation that all synonymous codons are used equally. Using equations 5 and 6, we replace this by a normalized Relative Synonymous Codon Usage of Species (RSCUS) value

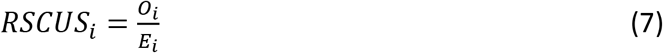

where *O_¿_* is the observed frequency with which amino acid *a* is encoded by codon *i*.

#### Controlling for Amino Acid Composition

Some amino acids may be more intrinsically prone to codon bias. We want a metric which quantifies effectiveness of selection (not amino acid frequency), so we re-weight CAIS for amino acid composition, to remove the effect of variation among species in amino acid frequencies.

Let *F_a_* be the frequency of amino acid *a* across the entire dataset of 118 vertebrate genomes, and *f_is_* the frequency of codon *i* in given species s. A candidate CAIS that controls for GC content but not for amino acid composition can be written as

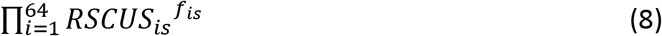

We want to re-weight *f_is_* on the basis of *F_a_* to ensure that differences in amino acid frequencies among species do not affect CAIS, while preserving relative codon frequencies for the same amino acid. We do this by solving for *a_as_* so that

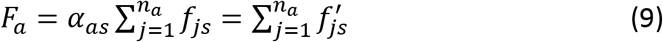

This gives us the amino acid frequency adjusted CAIS:

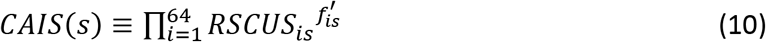

For convenient implementation in code, we used the following form:

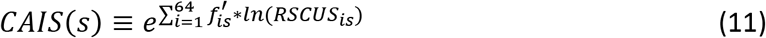

The *F_a_* values are available in a flatfile at https://github.com/cweibel2018/More_adapted_species_have_higher_SD/CAIS_ENC_calculation/Total_amino_acid_frequency_vertebrates.txt.

#### Novembre’s Effective Number of Codons (ENC) controlled for GC Content

Following from equations 5 and 6, the 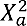 value representing the deviation of the frequencies of the codons for amino acid *a* from null expectations is

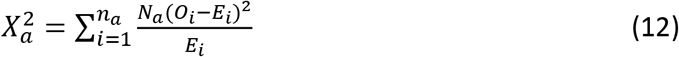

where *N_a_* is the total number of times that amino acid *a* appears. (Novembre 2002) defines the corrected “F value” of amino acid *a* as

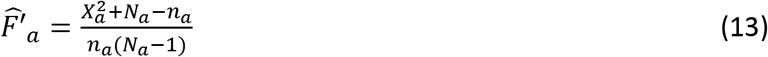

and the Effective Number of Codons as

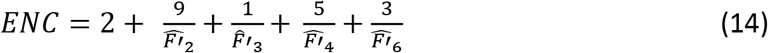

where each 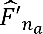 is the average of the “F values” for amino acids with *n_a_* synonymous codons. Past measures of ENC do not contain stop or start codons (Wright 1990; Novembre 2002; Fuglsang 2004), but as we expect variation in stop codon usage between species, and to facilitate more direct comparison with CAIS, we include stop codons as an “amino acid” and therefore amend (10) to

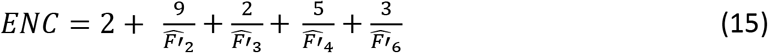

## Statistical Analysis

All statistical modelling was done in R 3.5.1. Scripts for calculating CAI and CAIS were written in Python 3.7. All code is available on https://github.com/cweibel2018/More_adapted_species_have_higher_SD.git, along with csv files containing CAIS values and ISD Effects on Species

### Linear model of species effect, controlling for Pfam identity

Linear models were implemented using the package lme4 (Bates et al, 2014). We used a mixed linear model to quantify the effect of species (fixed effect) on ISD with Pfam domain identity as a random effect, i.e. ISD ~ fixed(species identity) + random(Pfam identity). We ran three models of this form, from which we extracted species effects and standard errors for species effects on ISD for total, young, and old datasets. Note that running this model for all 118 species takes significant computational resources; our model using the total dataset required 144GB of RAM and ran for 8 CPU hours on 30 nodes of the UArizona HPC Ocelote server.

### Phylogenetic Independent Contrasts

Spurious phylogenetically confounded correlations can occur when closely related species share similar values of both metrics. One danger of such pseudoreplication is Simpson’s paradox, where there are negative slopes within taxonomic groups and a positive slope among them might combine to yield an overall positive slope. We avoid pseudoreplication by using Phylogenetic Independent Contrasts (PIC) (Felsenstein 1985) to assess correlation. PIC analysis was done using the R package “ape” (Paradis and Schliep 2019).

### Weighting linear models

PIC yield incommensurable units between analyses. To be able to compare the relationships of young domains to old, we used models of the form lm(species effect~CAIS, weights = (1/(Std._Error))^(2)), weighted by the standard errors of species effects. We extracted slopes of CAIS vs ISD and their standard errors.

## Acknowledgements

This work was supported by the National Institutes of Health (GM-104040), the John Templeton Foundation (60814), the Arnold and Mabel Beckman Foundation Scholars Program, the Western Alliance to Expand Student Opportunities (WAESO) Louis Stokes Alliance for Minority Participation (LSAMP) National Science Foundation (NSF) Cooperative Agreement (HRD-1101728), and UA/NASA Space Grant Undergraduate Research Internship program. We thank Luke Kosinski for helpful discussions and the University of Arizona Undergraduate Biology Research Program for training.

## Supplementary material

**Supplementary Figure 1:**
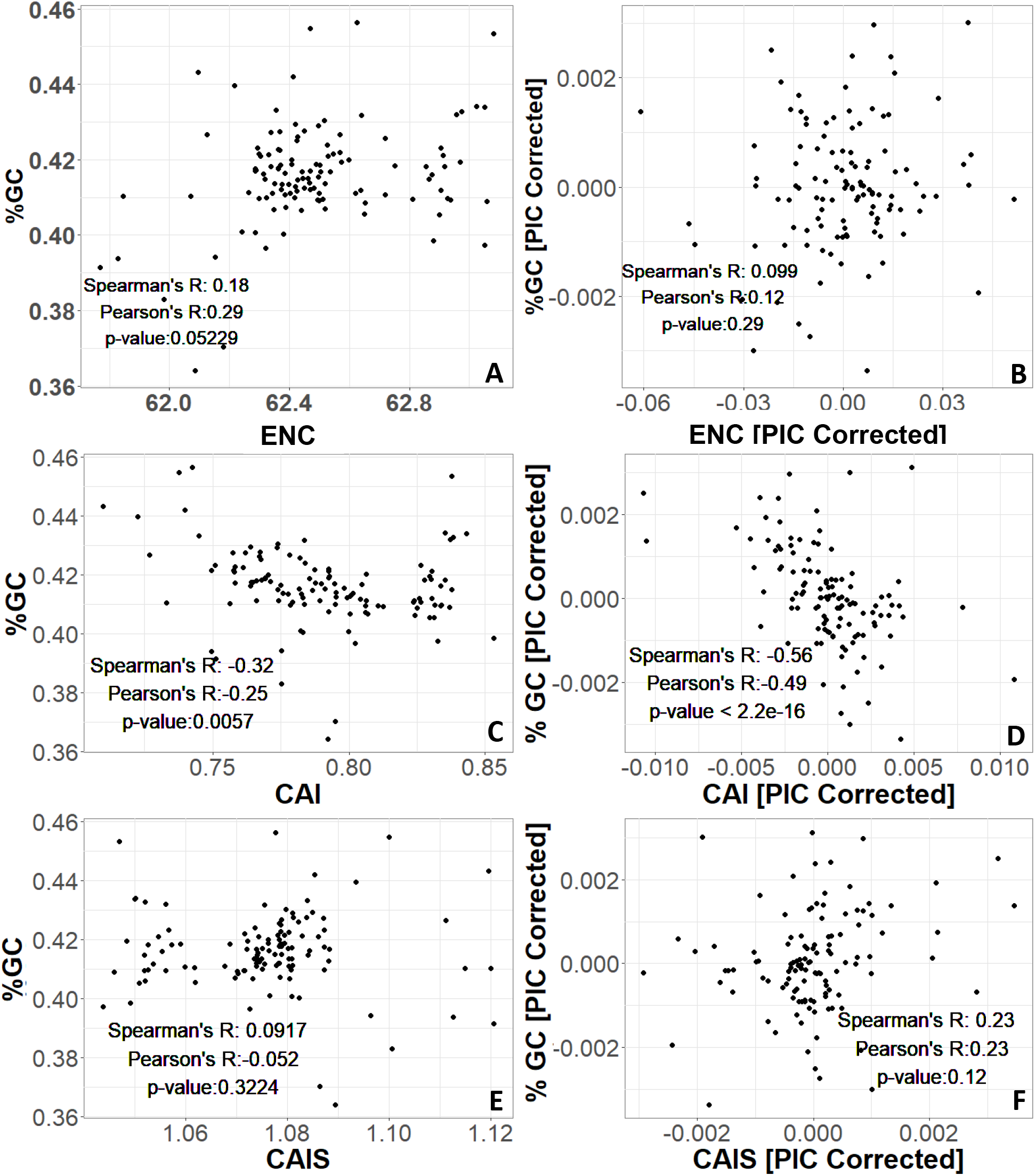
Effective Number of Codons (ENC) and Codon Adaptation Index of Species (CAIS) are uncorrelated with total genomic GC Content. Each datapoint is one of 118 vertebrate species with Complete intergenic genomic sequence available for %GC, and TimeTree divergence dates. Results are robust to controlling for phylogenetic confounding via Phylogenetic Independent Contrasts (PIC). P-values shown are for Spearman’s correlation.

**Supplementary Figure 2:**
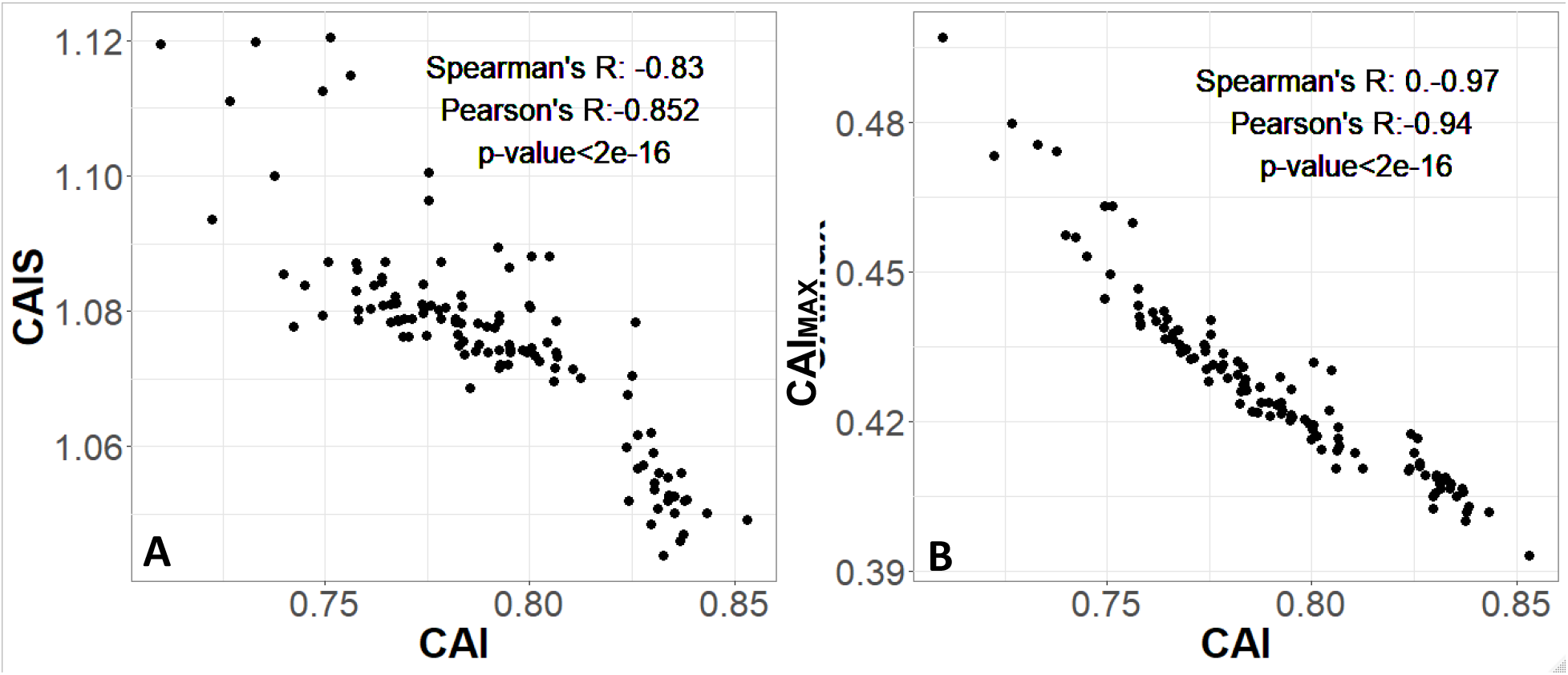
Codon Adaptation Index is not appropriate for species-wide effectiveness of selection measurements, because its value is driven by its normalizing denominator term. Each CAI value is averaged over an entire species’ genome. Each datapoint is one of 118 vertebrate species with Complete intergenic genomic sequence available for %GC, and TimeTree divergence dates. P-values shown are for Spearman’s correlation.

**Supplementary Figure 3:**
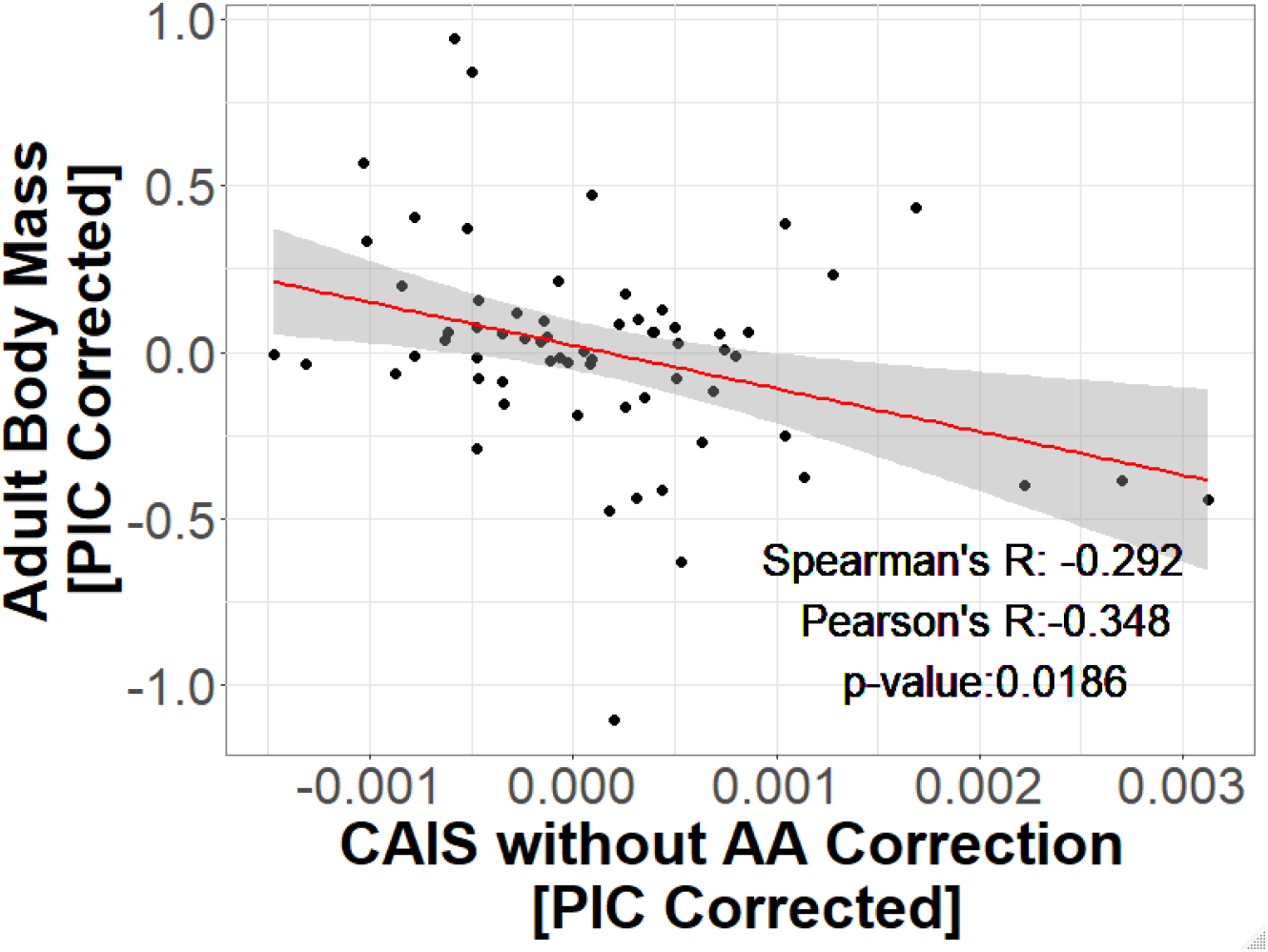
CAIS without correction for amino acid composition still reflects the expected relationship between effectiveness of selection and body. Body size data from PanTHERIA database, originally in log10(mass) in grams; data shown for 62 species in common between PANTHERIA and our own dataset of 118 vertebrate species with Complete intergenic genomic sequence available for %GC, and TimeTree divergence dates. P-value is shown for Spearman’s correlation. Red line shows unweighted lm(y~x) with grey region as 95% confidence interval.

**Supplementary Figure 4:**
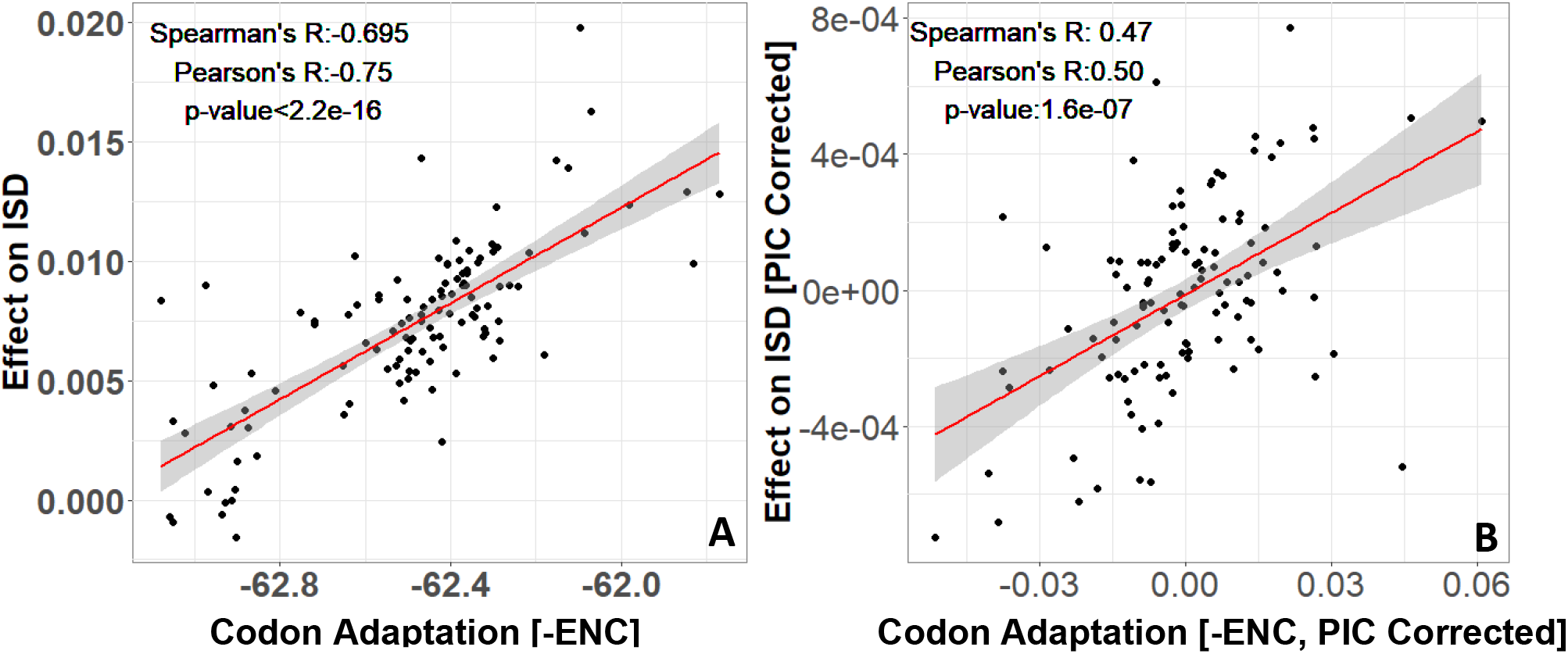
Our Figure 3 finding that more exquisitely adapted species have protein domains with higher ISD is confirmed by ENC. The same n=118 datapoints are shown as in Figure 3. P-values shown are for Spearman’s correlation. Red line shows unweighted lm(y~x) with grey region as 95% confidence interval.

**Supplementary Figure 5:**
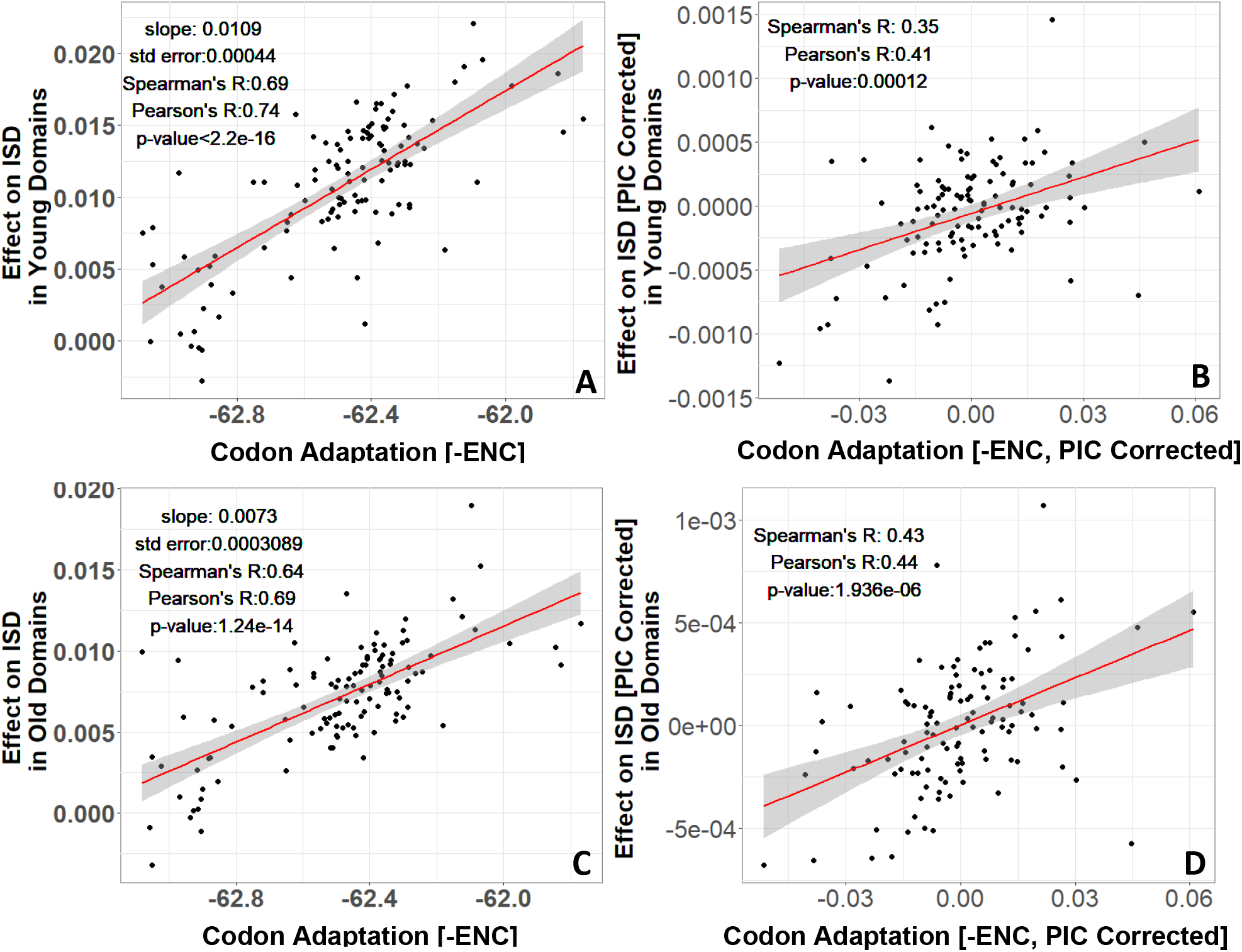
More exquisitely adapted species have higher ISD in both ancient and recent protein domains, as confirmed by ENC. Protein domains that emerged prior to LECA are identified as “old”, and protein domains that emerged after the divergence of animals and fungi from plants and found in vertebrates are identified as “young”. Age assignments are taken from (James et al. 2020). The same n=118 datapoints are shown as in Figure 2. P-values shown are for Spearman’s correlation. Red line shows lm(y~x),with grey region as 95% confidence interval, using a weighted model for non-PIC-corrected figures and unweighted for PIC-corrected figures. Weighted models make more accurate the comparison between young/old domains.

